# Molecular mechanism of β-arrestin-2 interaction with phosphatidylinositol 4,5-bisphosphate

**DOI:** 10.1101/2024.01.22.576757

**Authors:** Kiae Kim, Ka Young Chung

## Abstract

Phosphorylated residues of G protein-coupled receptors bind to the N-domain of arrestin, resulting in βXX release. This induces further allosteric conformational changes, such as polar core disruption, alteration of interdomain loops, and domain rotation, which transform arrestins into the active state. It is widely accepted that arrestin activation occurs by conformational changes propagated from the N-to the C-domain. However, recent studies have revealed that binding of phosphatidylinositol 4,5-bisphosphate (PIP_2_) to the C-domain transforms arrestins into an active state. In this study, we aimed to elucidate the mechanisms underlying PIP_2_-induced arrestin activation. We compared the conformational changes of β-arrestin-2 upon binding of PIP_2_ or phosphorylated C-tail peptide of vasopressin receptor type 2 using hydrogen/deuterium exchange mass spectrometry (HDX-MS). Introducing point mutations on the potential routes of the allosteric conformational changes and analyzing these mutant constructs with HDX-MS revealed that PIP_2_-binding at the C-domain affects the back loop, which destabilizes the gate loop and βXX to transform β-arrestin-2 into the pre-active state.

## Introduction

Arrestins are a protein family that regulate G protein-coupled receptor (GPCR) signaling, with four distinct members in mammals (arrestin 1–4) (Benovic *et al*, 1987; Lohse *et al*, 1990). Among these, arrestin-1 and –4 are expressed in the visual system, while arrestin-2 and –3 (β-arrestin-1 [βarr1] and 2) are found throughout various tissues (Lohse & Hoffmann, 2014). They interact with agonist-activated phosphorylated GPCRs, leading to receptor desensitization and internalization (Benovic *et al*., 1987). In addition, arrestins play a role in regulating other signaling pathways, including the mitogen-activated protein kinase signal transduction cascade (Coffa *et al*, 2011; Park *et al*, 2019; Perry-Hauser *et al*, 2022; Perry *et al*, 2019; Qu *et al*, 2021; Smith & Rajagopal, 2016; Srivastava *et al*, 2015). Understanding how arrestins are activated at the structural and molecular level is crucial for the development of drugs targeting GPCRs or other signaling cascades regulated by arrestins.

Various biophysical and biochemical techniques have been used to examine arrestin structures in both the basal and receptor-bound active states (Chen *et al*, 2023; Huang *et al*, 2020; Lee *et al*, 2020; Mayer *et al*, 2019; Park *et al*., 2019; Shukla *et al*, 2014; Staus *et al*, 2020; Yang *et al*, 2015; Yun *et al*, 2015; Zhou *et al*, 2017). These studies showed that arrestins consist of N– and C-domains, and two stable interactions stabilize the basal state (Figure 1A). The first is the interaction between the C-tail, more precisely βXX, and residues at the N-domain (Figure 1A, purple circle). The other region is the polar core formed by ionic interactions between residues within the gate loop, βIII, βX, and C-tail (Figure 1A, cyan circle). Binding of the phosphorylated residues of a GPCR (Figure 1B, green) at the N-domain, where βXX is located in the basal state, transforms arrestins into the active state via allosteric conformational changes. In the receptor-bound active state, βXX is released from the N-domain, the polar core is disrupted, the loops between the N– and C-domains alter their conformation, and the relative orientation between the N– and C-domains shifts (*i.e*., domain rotation) (Figure 1B). Although these conformational changes are the “canonical” changes of the receptor-activated arrestins, the degree of these changes can vary depending on the receptor types and phosphorylation patterns, which results in different active status of arrestins and different functional outcomes (Kaya *et al*, 2020; Latorraca *et al*, 2020; Maharana *et al*, 2023; Mayer *et al*., 2019; Yang *et al*., 2015; Zhou *et al*., 2017)

**Figure 1:**
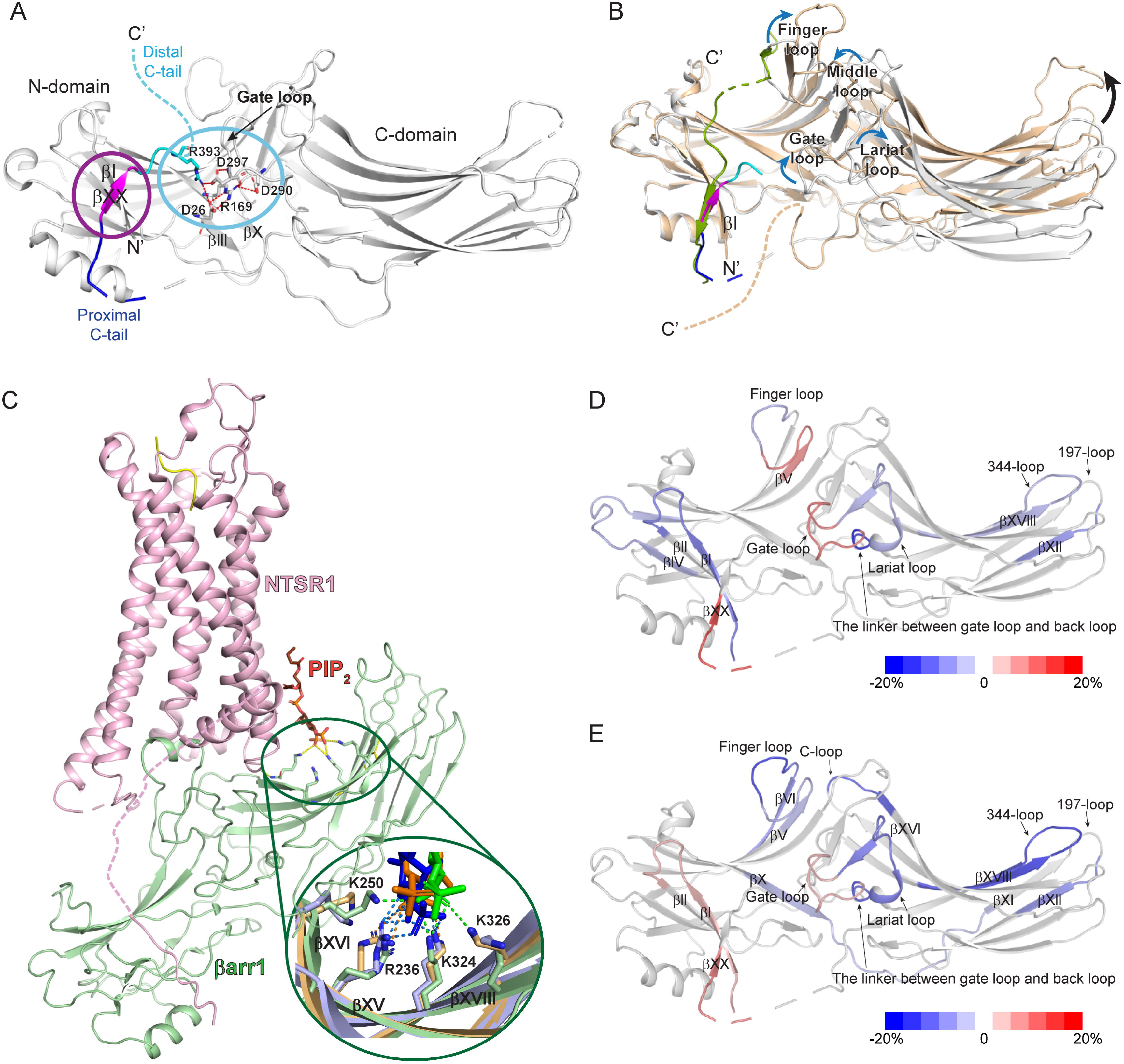
Structures of βarr in various states and HDX-MS profile changes upon the binding of V2Rpp or PIP_2_ to βarr2. A Structure of βarr1 in the basal state (PDB: 1G4R). The basal state βarr1 is colored gray with the C-terminus colored blue (proximal C-tail), magenta (βXX), and cyan (distal C-tail). Unresolved regions are indicated by dotted lines. The interaction between βXX and the residues in the N-domain is indicated in the purple circle, and the polar core is denoted in the cyan circle. Residues that are involved in the polar core formation are shown as sticks. B Comparison of the structure of βarr1 in basal (PDB: 1G4R) and V2Rpp-bound (PDB: 4JQI) states. V2Rpp-bound βarr1 is colored light orange and V2Rpp is colored green. The color-codes for basal state of βarr1 are same as those of panel A. The conformational changes of the loop regions are shown with blue arrows, and the domain rotation is indicated with a black arrow. C Structure of the NTSR1-βarr1 complex (PDB: 6UP7). NTSR1 is colored light pink, and βarr1 is colored light green. PIP_2_ is indicated with orange sticks. The residues that interact with PIP_2_ are shown as sticks. In the enlarged green circle, various modes of interaction between βarr1 and PIP_2_ are shown; PIP_2_ in the NTSR1-βarr1 complex (PDB: 6UP7) is colored orange, the interacting residues in βarr1 are colored light orange, and the ionic interactions between PIP_2_ and βarr1 are shown as green dotted lines; PIP_2_ in the GCGR1-βarr1 complexes (PDB: 8JRU and 8JRV) is colored blue or green, the interacting residues in βarr1 are colored light blue or light green, and the ionic interactions between PIP_2_ and βarr1 are shown as blue or orange dotted lines respectively. D HDX-MS profile comparison between the apo and V2Rpp-bound βarr2. The HDX level differences (*i.e.*, HDX levels of apo βarr2 – HDX levels of V2Rpp-bound βarr2) are color-coded on the basal state structure of βarr2 (PDB: 3P2D). Results were derived from three independent experiments. E HDX-MS profile comparison between the basal and PIP_2_-bound βarr2. The HDX level differences (*i.e.*, HDX levels of apo βarr2 – HDX levels of PIP_2_-bound βarr2) are color-coded on the basal state structure of βarr2 (PDB: 3P2D). Results were derived from three independent experiments.

Recently, phosphatidylinositol 4,5-bisphosphate (PIP_2_) was reported to be involved in arrestin activation. Arrestins can become “catalytically activated” (*i.e.*, active without receptor binding) with the assistance of PIP_2_ (Eichel *et al*, 2018). A subsequent study proposed that PIP_2_-binding is necessary for certain GPCR-arrestin interactions and that PIP_2_ promotes βarr activation (Janetzko *et al*, 2022). Notably, the cryo-electron microscopy structures of the neurotensin receptor 1 (NTSR1)-βarr1 and glucagon receptor (GCGR)-βarr1 complexes showed that PIP_2_ is bound at the C-domain of βarr1 (Figure 1C) (Chen *et al*., 2023; Huang *et al*., 2020). However, the precise structural mechanism by which PIP_2_ promotes arrestin activation or stabilizes its interaction with certain GPCRs remains to be elucidated.

In this study, we investigated the structural mechanisms underlying PIP_2_-induced arrestin activation. While we were preparing this manuscript, Zhai *et al*. (2023) published a paper suggesting a potential structural mechanism of βarr activation upon PIP_2_-binding by using ^19^F-NMR. The authors analyzed specific residues labeled with ^19^F. In contrast, here, we used hydrogen-deuterium exchange mass spectrometry (HDX-MS) to analyze the conformational changes of the whole protein. HDX-MS monitors the exchange between the amide hydrogen in the protein and deuterium in the solvent, which provides information about the conformational dynamics of the protein (Bai *et al*, 1993; Mayne, 2016), and HDX-MS has been successfully used to analyze the conformational dynamics of βarrs in various activation states (Kim *et al*, 2015; Min *et al*, 2020; Yun *et al*., 2015). In this study, we compared the conformational dynamics of the PIP_2_-induced and the phosphorylated C-tail peptide of the vasopressin receptor type 2 (V2Rpp)-induced active status of βarr2. We further investigated the activation mechanism by introducing mutations and analyzing the structural changes of the mutant constructs upon PIP_2_-binding.

## Results

### Conformational changes of **β**arr2 upon PIP_2_-binding

To investigate PIP_2_-induced conformational changes of βarr2, purified βarr2 was incubated with water-soluble PIP_2_ as described in the Materials and Methods. Subsequently, deuterium exchange was initiated on ice for various durations (10, 100, 1000, and 10,000 seconds). The peptic peptides used for the HDX-MS analyses are shown in Figure EV1, and the HDX-MS data analyzed in the present study are summarized in Dataset EV1. For comparative analysis, we also examined the changes in HDX levels upon V2Rpp-binding because V2Rpp is a well-established model system for understanding βarr interactions with phosphorylated receptor C-tails (Figure 1B) (Latorraca *et al*., 2020; Mayer *et al*., 2019; Shukla *et al*, 2013; Yang *et al*., 2015).

When the HDX levels were compared between the V2Rpp-bound βarr2 or βarr2 alone, HDX differences were observed at various regions. HDX levels of the V2Rpp-bound βarr2 were higher in the N-terminal part of the finger loop, gate loop, and proximal C-tail through βXX (Figure 1D and EV2, peptides 62 – 69, 292 – 302, and 382 – 389). HDX levels were lower in a few regions within the N-domain (βI, βIV through βV, and C-terminal part of the finger loop; peptides 1 – 19, 41 – 55, and 70 – 69), domain interfaces (the lariat loop and the linker between the gate loop and back loop; peptides 281 – 291 and 303 – 306), and a few regions within the C-domain (197-loop and βXVIII; peptides 195 – 201 and 324 – 338) (Figure 1D and EV2).

The HDX-MS analyses data well-reflected the V2Rpp-induced conformational changes of βarr2. V2Rpp (Figure 1B, green) interacts at the N-domain groove and near βI. Thus, lower HDX levels of the V2Rpp-bound βarr2 at the N-domain (specifically, βI and C-terminal part of the finger loop) probe the V2Rpp-binding in these regions. Additionally, V2Rpp-binding causes higher HDX levels in the gate loop and proximal C-tail through βXX, indicating conformational changes resulting from βXX release and polar core disruption. The changes in the levels of HDX at the domain interfaces suggests conformational changes in the loop regions at the domain interfaces and/or domain rotation upon V2Rpp-binding. Changes in HDX levels in the C-domain may reflect long-range allosteric conformational changes transmitted from the N-domain V2Rpp-binding site.

As HDX-MS analysis effectively probed the V2Rpp-induced activation of βarr2, we sought to analyze the conformational changes of βarr2 upon PIP_2_-binding. Based on the NTSR1-βarr1 and GCGR-βarr1 complex structures, PIP_2_ can interact with positively charged residues at βXV (R236 in the βarr1 sequence), βXVI (K250 in the βarr1 sequence), and βXVIII (K324 and K326 in the βarr1 sequence) (Figure 1C, inlet). The HDX-MS analysis revealed that the HDX levels at βXVIII become lower upon coincubation with PIP_2_ (Figure 1E and EV2, peptide 324 – 338), implying the binding of PIP_2_ to purified βarr2. However, the HDX levels of the peptides covering βXV (Figure 1E and EV2, peptide 219 – 239) and βXVI (Figure 1E and EV2, peptide 251 – 258) were not affected. This may be due to two reasons. First, HDX monitors the buffer exposure of the amide hydrogen atoms at the peptide backbone. The binding of PIP_2_ at βXV and βXVI occurs through the charge–charge interaction mediated by the amino acid side chains. If this interaction had not affected the backbone amide hydrogens, we would not have been able to detect changes in HDX levels in these regions even after ligand binding. Second, the interacting residues may differ slightly between the receptor-bound (*i.e.*, PIP2-bound state shown in the NTSR1-βarr1 and GCGR-βarr1 complexes) and unbound states (*i.e.*, current study). Even in the receptor-bound states, PIP_2_ interacted differently between the NTSR1-bound and GCGR-bound states (Figure 1C, inlet).

Interestingly, we observed higher HDX levels at βI, gate loop, and proximal C-tail through βXX (Figure 1E and EV2, peptides 1 – 19, 292 – 302, and 382 – 389), which is the canonical feature of the βarr activation (*i.e.*, βXX release and polar core disruption) (Figure 1B) (Kim *et al*., 2015; Shukla *et al*., 2014; Yun *et al*., 2015). Of note, the HDX levels at the gate loop and proximal C-tail through βXX were higher in the V2Rpp-bound state than the PIP_2_-bound state (Figure EV2, peptides 292 – 302 and 382 – 389), which suggests that the PIP_2_-bound state is not as fully active as the V2Rpp-bound state. Thus, these results suggest that the binding of PIP_2_ destabilizes the gate loop and the interaction of βXX at the N-domain, which may transform βarr2 so that it is more amenable to be activated (*i.e.,* pre-active state). The higher HDX levels at βI might be due to increased conformational dynamics at βXX, which could not be observed in the V2Rpp-bound state because V2Rpp interacts at βI.

Altered HDX was also evident at the finger loop, βVI, βX through βXI, 197-loop, C-loop, lariat loop, and the linker between the gate loop and back loop (Figure 1E and EV2, peptides 62 – 69, 70 – 76, 75 – 81, 168 – 186, 195 – 201, 246 – 250, 281 – 291, and 303 – 306). Although most of these regions were also affected by V2Rpp-binding, the HDX-MS profiles at the finger loop and its extension (*i.e.*, βVI) (Figure 1E and EV2, peptides 62 – 69, 70 – 76, and 75 – 81) and the lariat and gate loops (Figure 1E and EV2, peptides 281 – 291 and 292 – 302) differed between the V2Rpp– and PIP_2_-bound states, suggesting that these regions adopt different conformations between V2Rpp– and PIP_2_-bound states. Furthermore, βX through βXI and the C-loop were affected by the binding of PIP_2_, but not by the binding of V2Rpp (Figure 1E and EV2, peptides 168 – 186 and 246 – 250).

### Distal C-tail of **β**arr2 is not involved in PIP_2_-induced activation

The evidence from the HDX-MS analysis suggests that the interaction of PIP_2_ at the C-domain affects the conformational dynamics of βI, gate loop, and βXX (Figure 1E) potentially through the allosteric transmission of the conformational changes from the C-domain to the gate loop and βXX. Thus, we sought to understand the routes for the allosteric conformational changes transmitted from the PIP_2_-binding sites to the gate loop or βXX.

The initial candidate was the distal C-tail (Figure 2A). High-resolution structures have not fully characterized the distal C-tail because it is often unresolved or truncated (Han *et al*, 2001; Hirsch *et al*, 1999; Zhan *et al*, 2011). Nonetheless, given that the truncation of the distal C-tail transforms βarrs into the pre-active state (Celver *et al*, 2002; Gurevich, 1998; Gurevich *et al*, 1997; Kovoor *et al*, 1999), it is reasonable to hypothesize that the binding of PIP_2_ perturbs the conformational dynamics of the distal C-tail to impact the activation status of βarrs. To test this hypothesis, we truncated the distal C-tail (βarr2_1-394) and examined HDX level changes upon the binding of PIP_2_. If the distal C-tail serves as the route for allosteric conformational changes, PIP_2_ should not affect HDX levels at the gate loop or βXX in βarr2_1-394.

**Figure 2:**
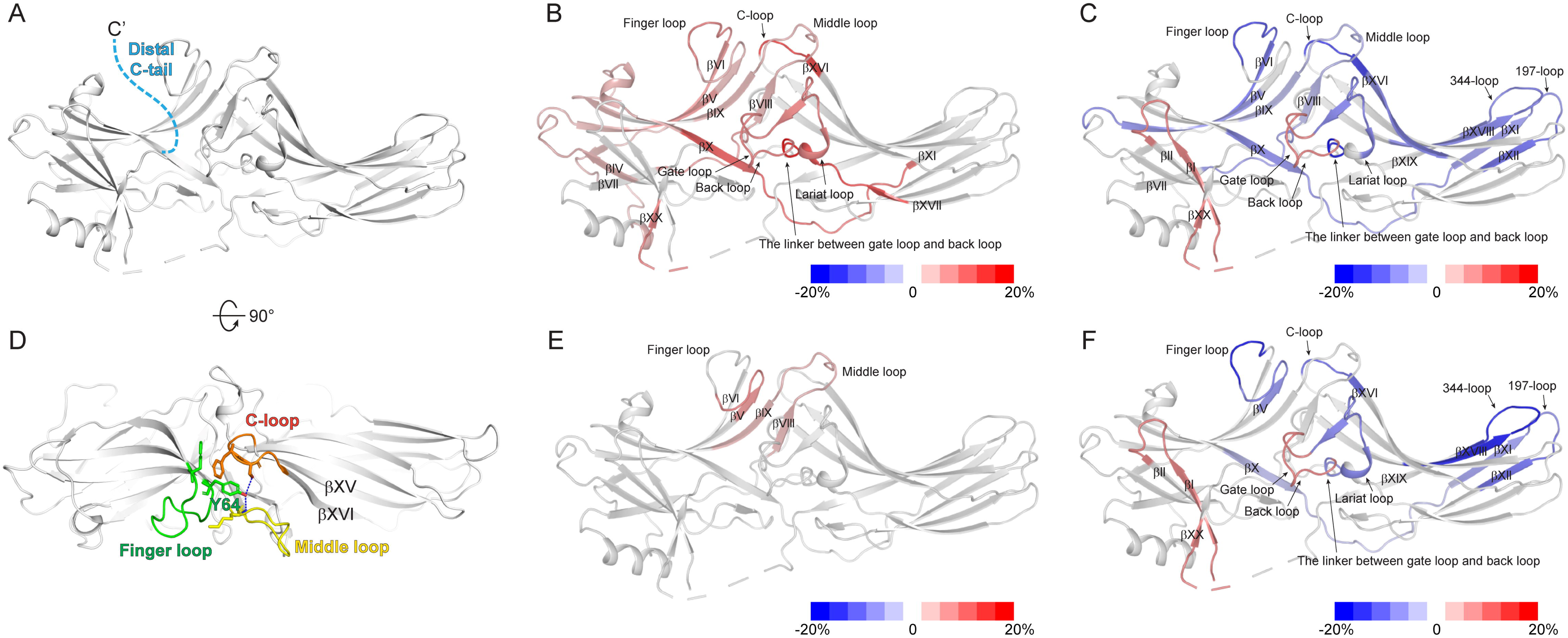
HDX-MS profile analysis of. β**-arrestin-2 (**β**arr2)_1-394 and Y64** A The truncated distal C-tail of βarr2 is colored light blue on the basal state structure of βarr2 (PDB: 3P2D). B HDX-MS profile comparison between the WT and βarr2_1-394. The HDX level differences (*i.e.*, HDX levels of WT βarr2 – HDX levels of βarr2_1-394) are color-coded on the basal state structure of βarr2 (PDB: 3P2D). Results were derived from three independent experiments. C HDX-MS profile comparison between apo and PIP_2_-bound βarr2_1-394. The HDX level differences (*i.e.*, HDX levels of apo βarr2_1-394 – HDX levels of PIP_2_-bound βarr2_1-394) are color-coded on the basal state structure of βarr2 (PDB: 3P2D). Results were derived from three independent experiments. D The top-view of the interaction between the finger, middle, and C-loops of βarr2 in the basal state (PDB: 3P2D). Y64 is indicated by a green stick. The finger, middle, and C-loops are green, yellow, and orange, respectively. E HDX-MS profiles of the WT and Y64A. HDX level differences (*i.e.*, HDX levels of WT βarr2 – HDX levels of Y64A) are color-coded based on the basal state structure of βarr2 (PDB: 3P2D). Results were derived from three independent experiments. F HDX-MS profile comparison of apo– and PIP_2_-bound Y64A. The HDX level differences (*i.e.*, HDX levels of apo Y64A – HDX levels of PIP_2_-bound Y64A) are color-coded based on the basal state structure of βarr2 (PDB: 3P2D). Results were derived from three independent experiments.

In the basal state, compared to the wild-type (WT), βarr2_1-394 exhibited higher HDX levels in numerous regions across the N– and C-domains (Figure 2B and EV3), indicating that the distal C-tail truncation yields βarr2 that is conformationally more dynamic. This increased conformational dynamics, especially at the gate loop and βXX, accounts for the pre-active state, as previously reported (Celver *et al*., 2002; Gurevich, 1998; Gurevich *et al*., 1997; Kovoor *et al*., 1999).

Upon the binding of PIP_2_, the HDX levels of βarr2_1-394 were altered in the regions similar to the WT (compare Figures 1E and 2C). Decreased HDX levels were detected at the PIP_2_-binding site (Figure 2C and EV3, peptide 324 – 338) and increased HDX levels were detected at βI, gate loop, and proximal C-tail through βXX (Figure 2C and EV3, peptides 1 – 19, 292 – 302, and 382– 389). These findings suggest that the binding of PIP_2_ can induce further activation of βarr2_1-394.

Other regions altered in the WT were also similarly affected (Figure 2C and EV3, peptides 70 – 76, 75 – 81, 168 – 186, 195 – 201, 246 – 250, 281 – 291, and 303 – 306). A few other regions where we did not observe HDX changes with PIP_2_-bound WT were also affected, but the HDX levels of these regions became similar to those of the WT (Figure EV3, peptides 50 – 64, 118 –127, 128 – 145, and 251 – 258). Additionally, we observed the decreased HDX levels at βXI (Figure 2C and EV3, peptide 187 – 194). Overall, the HDX profile changes of the PIP_2_-bound βarr2_1-394 (Figure 2C) were similar those of the PIP_2_-bound WT (Figure 1E). These results suggest that the distal C-tail is not the route for allosteric conformational changes from the PIP_2_-binding sites to the gate loop or βXX.

### Y64 in the finger loop is not involved in PIP_2_-induced activation

The finger, middle, and C-loops between the N– and C-domains undergo dramatic conformational changes upon activation (Figure 1B) and interact with the cytosolic core of the receptor (Figure 1C). In the basal state, the finger, middle, and C-loops form a designated structure through hydrophobic and polar interactions (Figure 2D). The binding of PIP_2_ altered HDX levels in the finger loop and C-loop (Figure 1E). Notably, the C-loop is located at the C-domain as an extension from the PIP_2_-binding sites (βXV and βXVI) (Figure 1C and 2D). Therefore, our second hypothesis was that allosteric conformational changes in the PIP_2_-binding sites are transmitted through interactions between the finger, middle, and C-loops. To test this hypothesis, we selected Y64 as a key residue. In the basal state, Y64 is located in a pocket formed by the finger-, middle-, and C-loops (Figure 2D), probably stabilizing the interactions between these three loops. Thus, the mutation of Y64 destabilizes the interactions between these three loops and breaks off the transmission route from the PIP_2_-binding sites.

In the PIP_2_-unbound state, the point mutation of Y64 to alanine (Y64A) altered HDX levels in the N-terminal half of the finger loop and middle loop compared to the those in the WT (Figure 2E and EV4, peptides 62 – 69 and 128 – 145), reflecting a disturbance of the conformation surrounding Y64, as expected.

Upon the binding of PIP_2_, Y64A displayed HDX changes in the regions similar to those of the WT (compare Figures 1E and 2F). HDX levels were decreased at the PIP_2_-binding site (Figure 2F and EV4, peptide 324 – 338) and increased at βI, gate loop, and proximal C-tail through βXX (Figure 2F and EV4, peptides 1 – 19, 292 – 302, and 382– 389). These findings suggest that the binding of PIP_2_ can induce pre-activation of Y64A. Other regions altered in the WT were also affected (Figure 2F and EV4, peptides 62 –69, 70 – 76, 168 – 186, 195 – 201, 246 – 250, 281 – 291, and 303 – 306). Additionally, we observed the decreased HDX levels at βXI (Figure 2F and EV4, peptides 187 – 194). These results suggest that our second hypothesis is incorrect; thus, the interactions between the finger, middle, and C-loops are not routes for allosteric conformational transmission.

### The lariat loop of **β**arr2 is involved in PIP_2_-induced activation

Because the distal C-tail and the interactions between the finger, middle, and C-loops do not serve as routes for allosteric conformational transmission, we sought other potential routes. After careful examination of the basal state structure and HDX-MS data of PIP_2_-bound βarr2, L280 in the lariat loop and E315 in the back loop were chosen as potential key residues (Figure 3).

**Figure 3:**
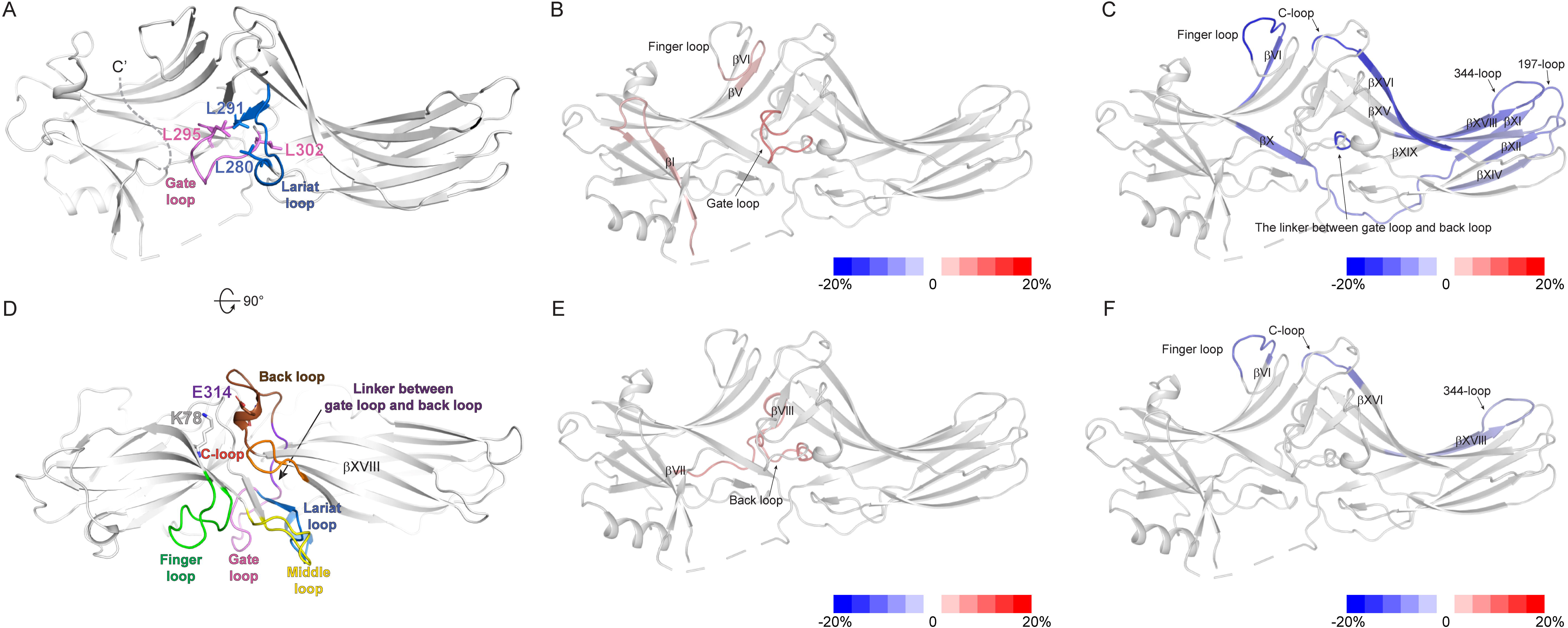
**HDX-MS profile analysis of L280 and E315** A Interaction between the lariat and gate loops in the basal state βarr2 (PDB: 3P2D). The lariat and gate loops are shown in blue and pink, respectively. The hydrophobic residues forming the interaction between the lariat and gate loops are shown as sticks. B HDX-MS profile comparison of the WT and L280G. The HDX level differences (*i.e.*, HDX levels of WT βarr2 – HDX levels of L280G) are color-coded on the basal state structure of βarr2 (PDB: 3P2D). Results were derived from three independent experiments. C HDX-MS profile comparison of apo and PIP_2_-bound L280G. The HDX level differences (*i.e.*, HDX levels of apo Y64A – HDX levels of PIP_2_-bound L280G) are color-coded on the basal state structure of βarr2 (PDB: 3P2D). Results were derived from three independent experiments. D Top-view of the structure of the basal state βarr2 (PDB: 3P2D) showing relative positions of the back, gate, and lariat loops, and the linker between the gate and back loops. E315 is shown as brown sticks, and K78 is shown as gray sticks. The finger, gate, lariat, middle, and C-loops, linker between the gate and back loops, and back loop are colored green, pink, blue, yellow, orange, violet, and brown, respectively. E HDX-MS profile comparison between the WT and E315A. The HDX level differences (*i.e.*, HDX levels of WT βarr2 – HDX levels of E315A) are color-coded on the basal state structure of βarr2 (PDB: 3P2D). Results were derived from three independent experiments. F HDX-MS profile comparison of apo and PIP_2_-bound E315A. The HDX level differences (*i.e.*, HDX levels of apo Y64A – HDX levels of PIP_2_-bound E315A) are color-coded on the basal state structure of βarr2 (PDB: 3P2D). Results were derived from three independent experiments.

HDX levels in the lariat loop were altered upon the binding of PIP_2_ (Figure 1E and EV2, peptide 281 – 291). In the basal state, L280 in the lariat loop faces the gate loop and forms hydrophobic interactions with L291, L295, and L302 (Figure 3A), which stabilizes the conformation of the gate and lariat loops. If allosteric conformational transmission is mediated through perturbation of the interaction of the lariat and gate loops, the L280 mutation would disrupt this route. To test this hypothesis, we mutated L280 to a glycine residue.

In the basal state, L280G showed altered HDX levels compared to those observed in the WT at the gate loop (Figure 3B and EV4, peptide 292 – 302), reflecting the altered conformation near the lariat and gate loop regions due to the mutation. We also observed altered HDX levels in L280G compared to those observed in the WT at βI and the N-terminal half of the finger loop (Figure 3B and EV4, peptides 1 – 19 and 62 – 69). The results suggest that perturbation of the interaction between the gate and the lariat loops could alter the conformational dynamics of remote regions, such as βI and finger loop.

Upon PIP_2_-binding to L280G, we observed decreased HDX levels at the PIP_2_-binding interface (Figure 3C and EV4, peptide 324 – 338). We also observed altered HDX at the regions similar to those of the WT (compare Figures 1E and 3C), such as the C-terminal part of the finger loop, βVI, βX through βXI, 197-loop, C-loop, and the linker between lariat loop and back loop (Figure 3C and EV4, peptides 70 – 76, 75 – 81, 168 – 186, 195 – 201, 246 – 250, and 303 – 306). Additionally, decreased HDX levels were evident at βXI, βXIV through βXV, and βXVI (Figure 3C and EV4, peptides 187 – 194, 219 – 239, and 251 – 258)

In contrast, changes in HDX levels for βI, the gate loop, and proximal C-tail through βXX were not evident upon the binding of PIP_2_ to L280G (Figure 3C and EV4, peptides 1 – 19, 292 – 302, and 382 – 389). These results suggest that the binding of PIP_2_ in L280G induces conformational changes in most regions similar to those of the WT but failed to transform it to the pre-active conformation (*i.e.*, disturbance of the gate loop and βXX). Therefore, the results of the HDX-MS analysis with L280G indicate that perturbation of the interaction of the lariat and gate loops is the route for the transmission of the conformational changes from the PIP_2_-binding site to βXX.

### The back loop of **β**arr2 is involved in PIP_2_-induced activation

Another potential route we examined was the back loop. Although the HDX-MS profiles of the back loop were not affected by the binding of PIP_2_, the neighboring C-loop and the linker between the lariat and back loops were altered (Figure 1E and EV2, peptides 246 – 250 and 303 – 306). Interestingly, the back loop is an extension of the PIP_2_-binding sites (βXVIII), located adjacent to the C-loop, and directly connected to the gate loop through the linker between the gate loop and the back loop (Figure 3D). Previous evidence suggested that in the basal state, E315 at the back loop occasionally forms salt bridge with K78 at βVI (Figure 3D) and that disruption of this interaction results in ligand-independent accumulation of βarr2 in the clathrin-coated endocytic structures (Eichel *et al*., 2018). Moreover, the back loop has been reported as a potential route for the conformational transition from PIP_2_-binding to βarr1 C-tail release (Zhai *et al*, 2023). Therefore, we further examined the role of the back loop in the βarr2 activation induced by the binding of PIP_2_.

To test this hypothesis, we mutated E315 to alanine. Compared to the HDX levels in the WT, E315A showed higher HDX levels at the back loop and its neighboring βVII/βVIII loop (Figure 3E and EV5, peptides 118 – 127 and 303 – 317) reflecting altered conformational dynamics of the back loop upon E315A mutation. As we did not observe HDX differences in other regions remote from the back loop, the results imply that the disruption of the interaction between E315 and K78 alters the local conformational dynamics but does not affect the overall conformational dynamics of βarr2.

Although E315A did not show appreciable differences in the conformational dynamics compared to the WT, the effects of the binding of PIP_2_ on E315A were dramatically different from those on the WT βarr2 (compare Figures 1E and 3F). Upon PIP_2_-binding to E315A, we observed a decrease in HDX levels at the PIP_2_-binding site (Figure 3F and EV5, peptide 324 – 338). However, we observed HDX-MS profile changes only within very limited regions, such as the C-terminal half of the finger loop and the C-loop (Figure 3F and EV5, peptides 70 – 76 and 246 – 250) but not other regions. These results suggest that in E315A the binding of PIP_2_ could alter the C-loop and its neighboring finger loop but fails to transform βarr2 to the pre-active state. Thus, we conclude that the conformational changes upon the binding of PIP_2_ is allosterically transmitted through the back loop to the gate loop and βXX.

## Discussion

No high-resolution structure of PIP_2_-bound βarr without receptors have been described. Previous studies with PIP_2_-bound β-arrestins monitored conformational changes of selected regions using fluorescence labeling or ^19^F-NMR at specific residues (Janetzko *et al*., 2022; Zhai *et al*., 2023). The current study comprehensively analyzed the conformational dynamics of the whole βarr2 using HDX-MS. The binding of PIP_2_ induced conformational changes near the PIP_2_-binding sites, loops between the N– and C-domains (i.e., finger, C-, lariat, and gate loops), βXX, and βI (Figure 1E).

The increased HDX levels at the gate loop and βXX reflect the active-like conformational status of PIP_2_-bound βarr2 because the release of βXX and disruption of the polar core are canonical features of the active arrestins (Kim *et al*., 2015; Yun *et al*., 2015). However, the extent of the activation was smaller than the V2Rpp-induced activation, suggesting that βXX is not fully released. This pre-active conformation of the PIP_2_-bound βarr2 has been previously proposed (Huang *et al*., 2020; Zhai *et al*., 2023).

However, the PIP_2_-binding sites are remote from these regions, and thus there must be allosteric conformational transmission from the PIP_2_-binding sites to the polar core and βXX. It has been suggested that the back loop is a potential route for transmitting structural changes triggered by the PIP_2_-binding to the polar core (Zhai *et al*., 2023). In this study, we provide further details on the allosteric conformational changes. Introducing mutations and HDX-MS analyses of the whole protein upon PIP_2_-binding revealed that the lariat and back loops, but not the finger loop or distal C-tail, are the routes for the allosteric conformational changes connecting the PIP_2_-binding to βarr2 activation.

Both the disruption of the interaction between the lariat and gate loops (L280G) or the alteration of the back loop (E315A) failed to activate βarr2 (Figure 3). However, the two mutants exhibited different conformational changes in other regions upon the binding of PIP_2_. In L280G, PIP_2_-binding could still induce conformational changes in almost all the regions, similar to the WT, except the gate loop, βXX, and βI (Figure 3C). This suggests that L280G could undergo conformational changes similar to the WT, but that these conformational changes would not be transferred to the gate loop or βXX when the interaction of the gate and lariat loops is disrupted. In contrast, in E315A, the binding of PIP_2_ induced conformational changes in limited regions (*i.e*., the C-loop and its neighboring finger loop; Figure 3F). The C-loop is an extension of the PIP_2_-binding sites (Figure 1C). Thus, in E315A, the binding of PIP_2_ affected the C-loop and its neighboring finger loop but could not induce further allosteric conformational changes.

These results suggest that the binding of PIP_2_ (Figure 4B, step 1) affects the conformational dynamics of the back loop (Figure 4B, step 2), which proceeds to the lariat loop/gate loop region (Figure 4B, step 4). The transmission from the back loop to the lariat loop/gate loop region were not through the finger loop or middle loop regions, as the Y64A mutant did not affect PIP_2_-induced βarr2 activation. Rather, the linker between the gate and back loops might be the route from the back loop to the lariat loop/gate loop region (Figure 4B, step 3), because we observed HDX profile changes in this region in the WT and other mutant constructs but not in E315A.

**Figure 4:**
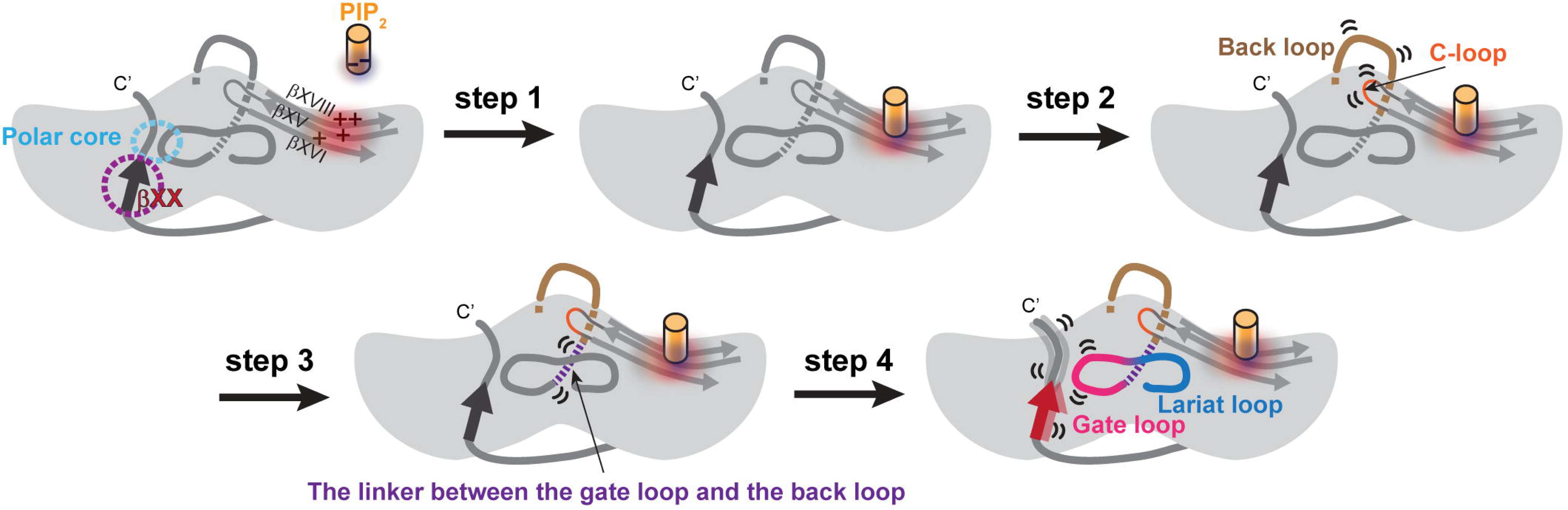
Proposed molecular mechanism of. β**-arrestin-2 pre-activation upon PIP_2_-binding.** Summary cartoon illustrating the proposed βarr2 pre-activation mechanism upon PIP_2_-binding.

It has long been believed that the interaction between phosphorylated residues of GPCRs in the N-domain is the key process for arrestin activation (Edward Zhou *et al*, 2019; Gusach *et al*, 2023; Hilger *et al*, 2018; Maharana *et al*, 2022; Seyedabadi *et al*, 2021; Wisler *et al*, 2014; Zhao *et al*, 2017). However, now it is emerging that arrestin activation can be achieved through various processes. Previously, the βarr2 structure bound to inositol hexaphosphate (IP_6_) revealed that IP_6_-binding at the phosphate sensor within the N-domain induces βXX release, activates βarr2, and triggers further downstream signal transduction (Chen *et al*, 2017). Recent studies suggested that PIP_2_ binds to positively charged residues in the C-domain to activate arrestins (Janetzko *et al*., 2022; Zhai *et al*., 2023), and here we further propose the structural mechanism of PIP_2_-induced arrestin pre-activation. As it is evident that arrestins can be activated via various routes, it is needed to investigate the structural differences and the functional consequences of the different active states of arrestins.

## Materials and Methods

### **β**-arrestin-2 expression and purification

All protein constructs were cloned into the pET28a vector and transformed into *Escherichia coli* BL21 (DE3). Expression and purification were performed as previously described (Park *et al*., 2019). Briefly, WT rat β-arrestin-2 and the mutants were grown in LB broth medium at 37 °C until the optical density at 600 nm reached 0.4 – 0.6. The bacteria were then induced with 0.03 mM IPTG for 24 h at 16 °C. Proteins were purified using Ni-IDA resins and size-exclusion chromatography.

### Hydrogen/deuterium exchange

β-arrestin-2 protein samples at a final concentration of 50 μM in 20 mM HEPES pH 7.4, 150 mM NaCl, and 100 μM Tris [2-carboxyethyl] phosphine hydrochloride were co-incubated with 500 μM V2Rpp and 150 μM PIP_2_ for 1 h at room temperature. HDX was performed by mixing 2 μl of protein (50 μM) with 28 μl of D_2_O buffer (20 mM HEPES pH 7.4, 150 mM NaCl, 100 μM Tris [2-carboxyethyl] phosphine hydrochloride, and 10% glycerol in D_2_O) and incubating the mixture for 10, 100, 1000, and 10,000 seconds on ice. At the indicated time points, the reaction was quenched by adding 30 μl of ice-cold quench buffer (60 mM NaH_2_PO_4_ pH 2.01 and 10% glycerol) and snap-frozen on dry ice. Identical procedures were conducted for nondeuterated samples using a H_2_O buffer comprising 20 mM HEPES, pH 7.4, 150 mM NaCl, and 10% glycerol in H_2_O.

### HDX-MS

HDX-MS and data analyses were conducted as previously described (Du *et al*, 2019; Qu *et al*., 2021). Briefly, the quenched samples underwent online digestion by passage through an immobilized pepsin column (2.1 × 30 mm; Life Technologies, Carlsbad, CA, USA). The digested peptide fragments were collected on a C18 VanGuard trap column (1.7 mm × 30 mm; Waters, Milford, MA, USA), followed by ultra-pressure liquid chromatography using an ACQUITY UPLC C18 column (1.7 mm, 1.0 mm × 100 mm; Waters). All settings and conditions for the system, such as voltage, temperature, collision energy, and lockspray, were as previously reported (Du *et al*., 2019; Qu *et al*., 2021). Peptic peptides from nondeuterated samples were identified using ProteinLynx Global Server 2.4 (Waters). To process HDX-MS data, the amount of deuterium in each peptide was determined by measuring the centroid of the isotopic distribution using DynamX 3.0 (Waters).

### Statistical analysis

For HDX-MS analysis, mass differences >0.22 Da and 2% were considered significant. Student’s t-test was used to determine the statistically significant differences between individual time points using GraphPad Prism software (GraphPad, San Diego, CA, USA). Statistical significance was set at p < 0.05.

## Acknowledgments

This work was supported by grants from the National Research Foundation of Korea, funded by the Korean government (NRF-2021R1A2C3003518 and NRF-2019R1A5A2027340 to K.Y.C.).

## Author Contributions

**Kiae Kim:** Investigation; formal analysis; writing the original draft. **Ka Young Chung:** Conceptualization, funding acquisition, project administration, investigation, formal analysis, writing and original draft.

## Data availability

This study did not include data from external repositories. All the data supporting the findings of this study have been included in the manuscript.

**Expanded View** for this article is available online.

