## Supplementary material for "Molecular mechanism of β-arrestin-2 interaction with phosphatidylinositol 4,5-bisphosphate": Fig EV

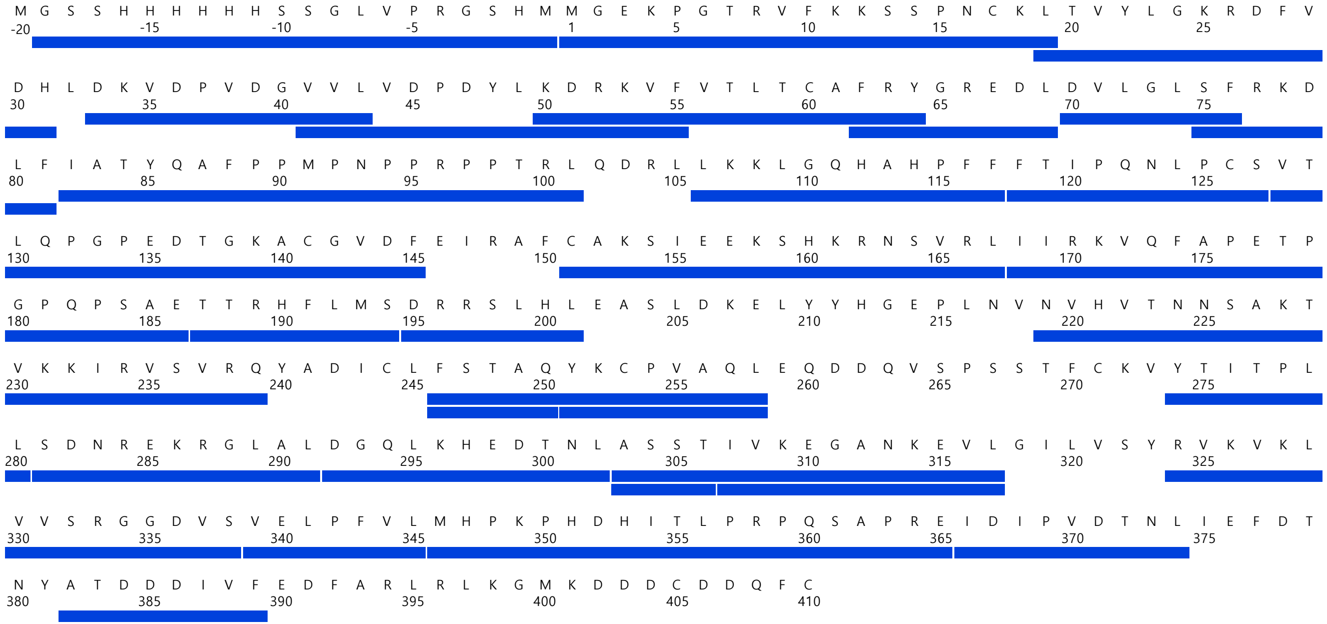


Figure EV1. Sequence coverage map of wild-type β-arrestin-2 (βarr2).
The blue bars indicate analyzed peptic peptides.

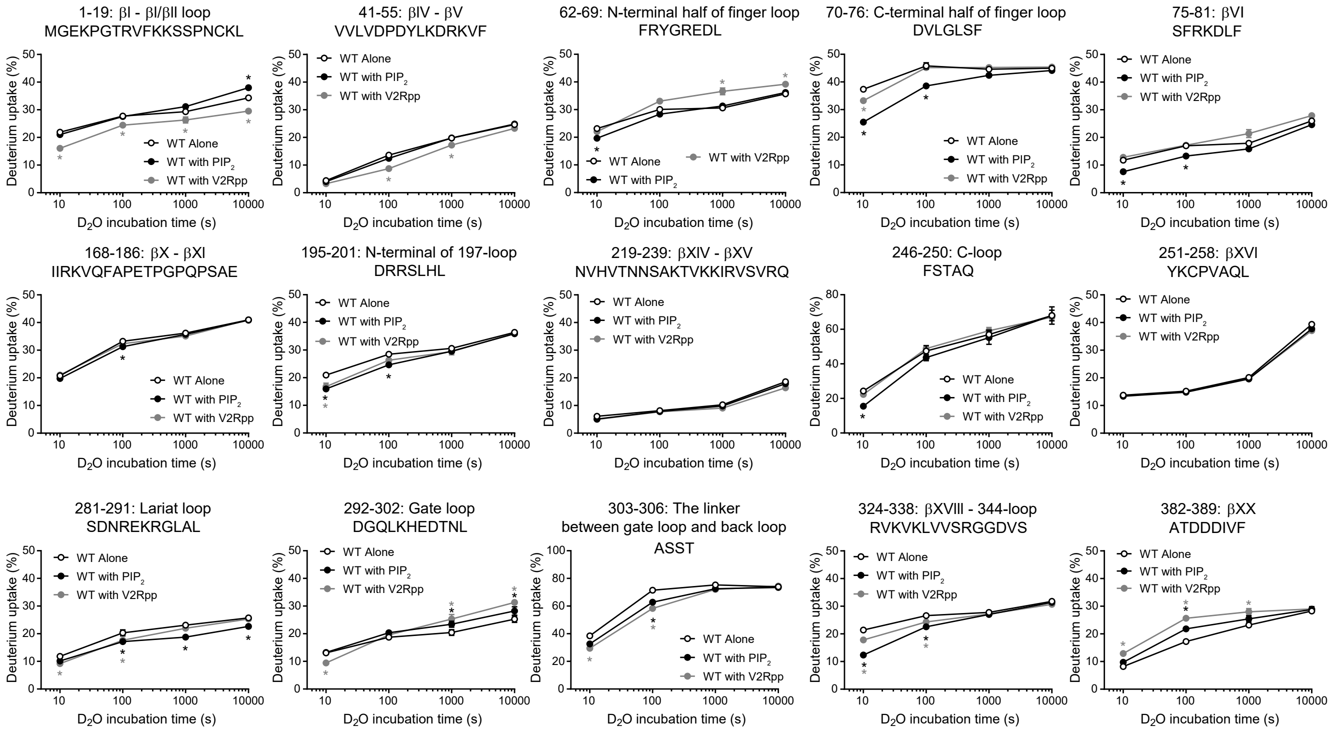


Figure EV2. Deuterium uptake plots of selective peptides of WT βarr2 with or without V2Rpp or $PIP_2$ co-incubtion. Results were derived from three independent experiments. The statistical significance of the differences was determined using Student's t-test (*$p < 0.05$). Data are presented as mean ± standard error of the mean. Black or grey * indicates statistically significant difference between apo WT βarr2 and $PIP_2$-bound WT βarr2 or V2Rpp-bound WT βarr2, respectively.

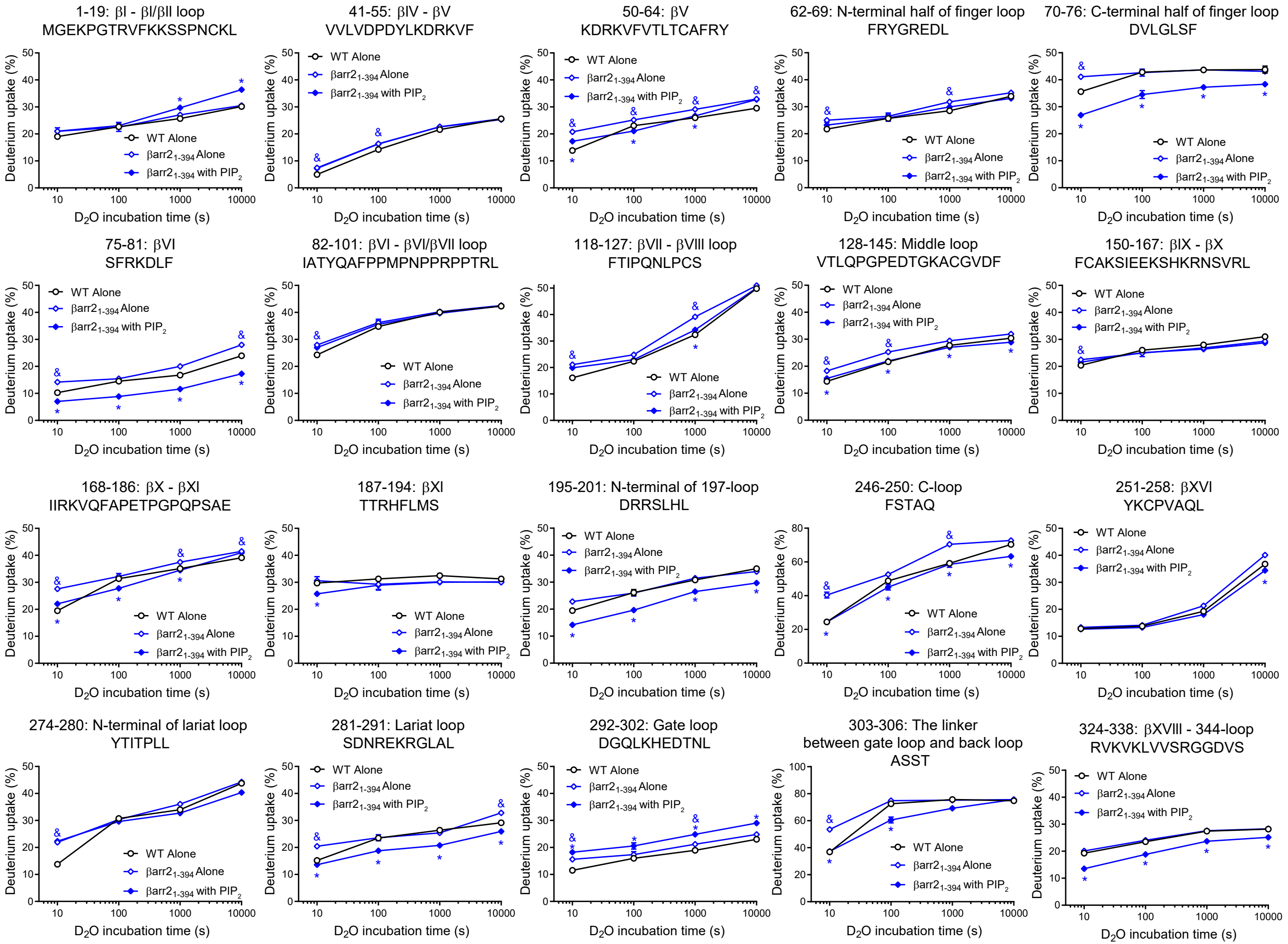


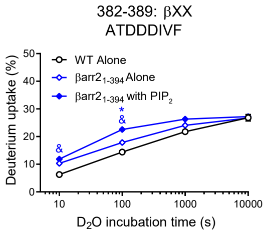


Figure EV3. Deuterium uptake plots of selective peptides of apo WT βarr2 and βarr2_1-394 with or without $PIP_2$ co-incubation

Results were derived from three independent experiments. The statistical significance of the differences was determined using Student's t-test (&,* $p < 0.05$). & indicates statistically significant difference between apo WT βarr2 and apo βarr2_1-394. * indicates statistically significant difference between apo and $PIP_2$-bound βarr2_1-394. Data are presented as mean ± standard error of the mean.

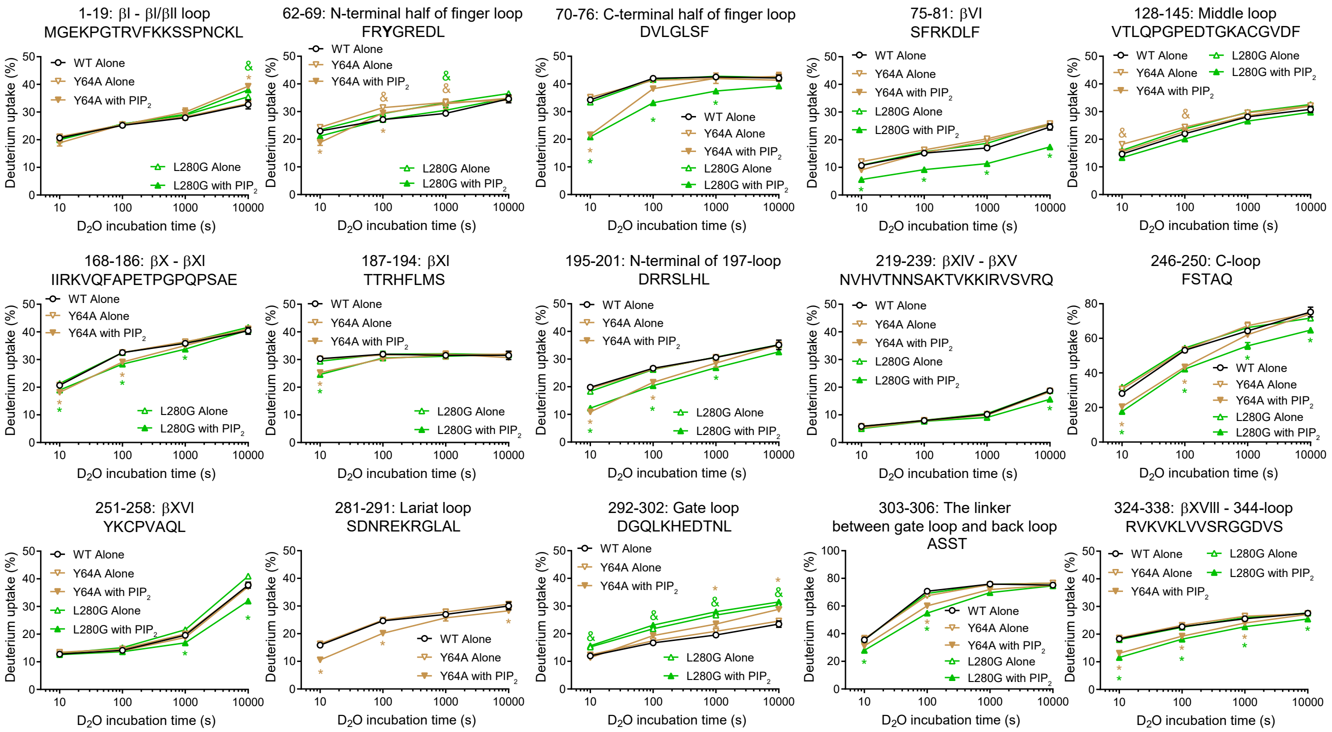


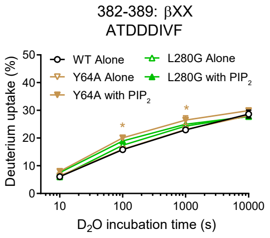


Figure EV4. Deuterium uptake plots of selective peptides of apo WT βarr2 and Y64A and L280G with or without $PIP_2$ co-incubation

Results were derived from three independent experiments. The statistical significance of the differences was determined using Student's t-test (&,* $p < 0.05$).

Yellow orcher or green & indicates statistically significant difference between apo WT β arr2 and apo Y64A or L280G, repsectively. Yellow orcher or green * indicates statistically significant difference between apo and $PIP_2$-bound Y64A or L280G, respectively. Data are presented as mean ± standard error of the mean.

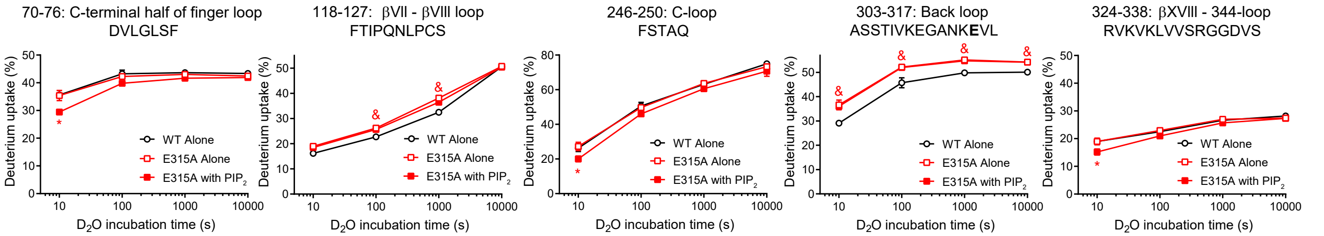


Figure EV5. Deuterium uptake plots of selective peptides of apo WT βarr2 and E315A with or without $PIP_2$ co-incubation

Results were derived from three independent experiments. The statistical significance of the differences was determined using Student's t-test (&,* $p < 0.05$). & indicates statistically significant difference between apo WT βarr2 and apo E315A. * indicates statistically significant difference between apo and $PIP_2$-bound E315A. Data are presented as mean ± standard error of the mean.
